# Hydration-Controlled Proton Transport in Respiratory Complex I

**DOI:** 10.64898/2026.01.29.702666

**Authors:** Jong Ho Choi, Gregory A. Voth

**Affiliations:** Department of Chemistry, Chicago Center for Theoretical Chemistry, James Franck Institute, and Institute for Biophysical Dynamics, The University of Chicago, Chicago, Illinois 60637

## Abstract

Proton pumping by respiratory Complex I is one essential element for generating the proton motive force that drives ATP synthesis in mitochondria. Although it is understood that electrons from NADH reduce ubiquinone at the peripheral arm and that four protons are transferred in the membrane domain, the mechanism by which this redox reaction initiates proton translocation remains unclear. A lateral pathway linking the quinone binding site to the membrane domain via ND1, ND3, and ND4L subunits has been proposed as the initial path of an excess proton. However, in experimental structures this region lacks a continuous water network between D66_ND3_ and E34_ND4L_, resulting in a hydration bottleneck that may regulate proton transfer. Using multiscale reactive molecular dynamics (MS-RMD) and a water wire connectivity metric, we directly simulate proton transport through this region as coupled the the hydration by water molecules. Our results reveal that proton transfer is thermodynamically feasible when transient hydration aligns with the presence of an excess proton, revealing the strong coupling between hydration and proton (PT) in this region of Complex I. These findings support a model where proton injection enhances local hydration, dynamically opening the pathway for proton transfer and regulating the onset of proton pumping in Complex I.

## INTRODUCTION

Respiratory complex I (NADH:ubiquinone oxidoreductase) is the largest enzyme in the electron transport chain (ETC).^1-5^ During its catalytic cycle, two electrons from NADH oxidation are transferred through a chain of Fe–S clusters and reduce ubiquinone, and the free energy released in this redox reaction is harnessed to translocate four protons across from N-side to P-side, contributing to the proton motive force that ultimately drives ATP synthesis.^5-11^ Complex I is a massive protein cluster, approximately 200 Å in length, composed of a hydrophilic peripheral arm and a membrane domain (Figure 1). A central question regarding this enzyme is how electron transfer and redox reaction that occurs in the peripheral arm is coupled to proton pumping in the membrane domain.

**Figure 1.**
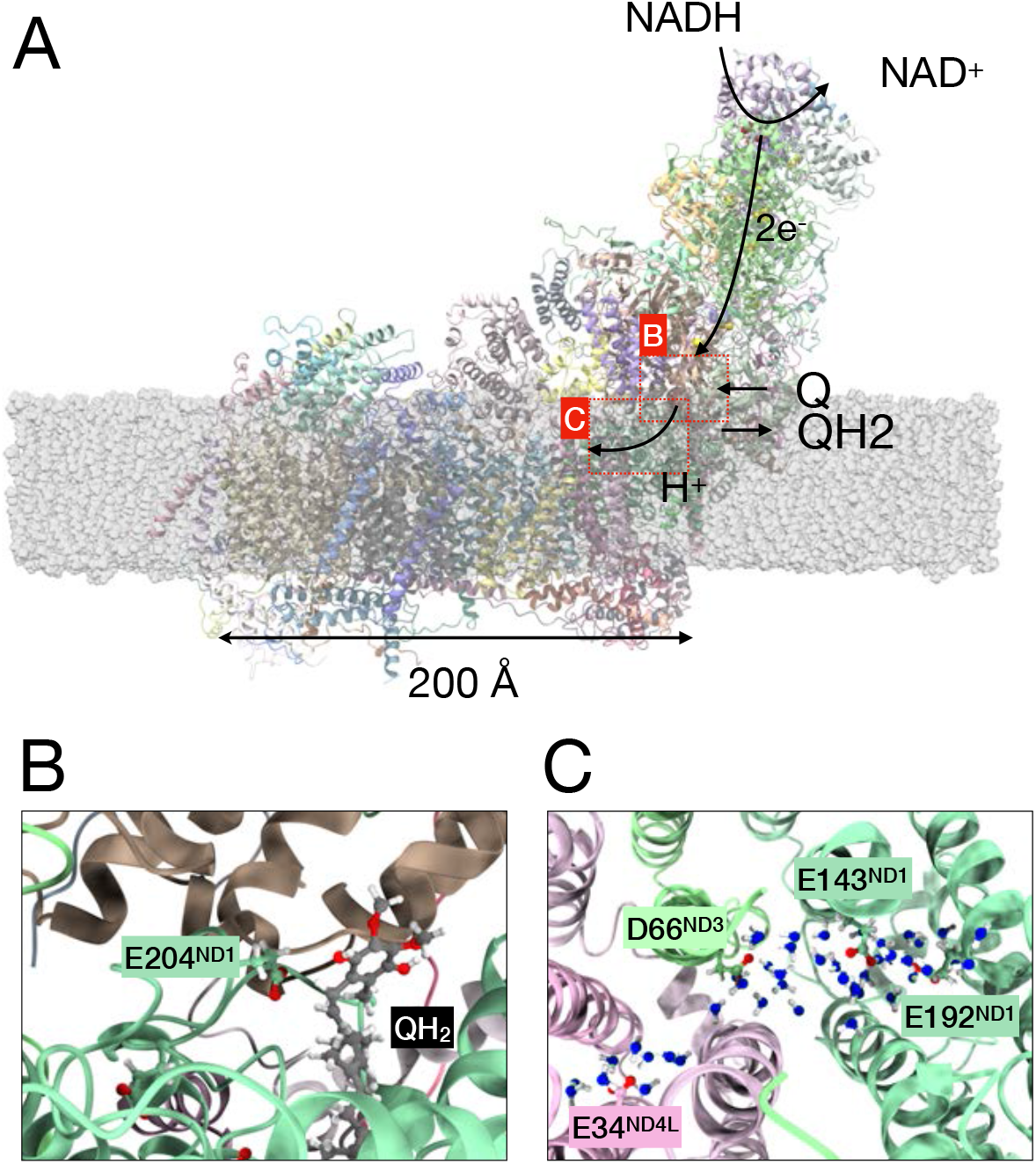
(A) The structure and working mechanism of respiratory Complex I. (B) The configuration of ubiquinol (QH_2_) and the entrance of E-channel. (C) The structure of water wire inside the PT channel through E192_ND1_ to E34_ND4L_. The oxygens of water molecules are represented as blue for easier comparison to the oxygens of GLU and ASP.

Ubiquinone reduction at the Q-binding site is proposed to trigger proton injection into a hydrophilic cavity within subunit ND1, commonly referred to as the E-channel.^9-17^ In recently suggested mechanistic models, after the reduction of ubiquinone, the semiquinone migrates toward the entrance of the E-channel, and this movement is accompanied by the uptake of protons from the N-side, which enter the E-channel and initiate proton pumping.^4, 7, 10^ The injected proton is believed to travel through the E-channel and the subsequent lateral path comprising the ND1, ND3, and ND4L subunits, which connects the Q-binding cavity to the membrane domain.^18-20^ This route is thus proposed to serve as a unique coupling interface between redox reactions and proton pumping in Complex I.

The ND1–ND3–ND4L segment represents the initial entry point of protons into the membrane domain and is structurally distinct from the other antiporter-like subunits (ND2, ND4, ND5) in the membrane domain which also transport protons.^4, 12, 13, 17, 21^ While the antiporter-like subunits exhibit relatively well-defined water pathways connecting the N-side to the P-side, structural studies have not resolved a continuous water network connecting ND4L to the P-side.^18, 22, 23^ Instead, a lateral water chain linking ND4L to ND2 has been observed, suggesting that protons reaching ND4L may diffuse horizontally and ultimately exit through adjacent subunits.^17, 18, 23^ This inter-subunit connectivity raises the possibility that proton translocation along the ND1–ND4L route initiates proton pumping in the following subunits of the membrane domain. Moreover, the short segment between D66_ND3_ and E34_ND4L_ has been consistently identified as a relatively dry region in both experimental^12, 20, 24-27^ and standard (non-reactive) molecular dynamics studies.^16, 18, 20^ This hydration bottleneck is hypothesized to be a regulatory gate, controlling proton transfer and, by extension, regulating the overall mechanism of proton pumping in Complex I.

Several previous simulation studies have investigated hydration patterns and possible proton transfer pathways in the ND1–ND4L region using classical molecular dynamics and Quantum Mechanics/Molecular Mechanics (QM/MM) methods.^12, 16-18, 20, 28^ These efforts typically assessed hydration by counting the number of water molecules within the pathway. While such metrics offer useful structural insight, they do not quantify the continuity of the water network or capture the dynamic role of hydration in proton translocation.^29^ Moreover, because proton transport (PT) involves bond breaking and formation as an excess proton shuttles between water molecules and (at times) protonatable amino acids, it cannot be modeled with traditional non-reactive or “classical” MD simulations. Although the QM/MM method can describe bonds breaking and forming, because of its demanding computational cost, QM/MM simulations are limited to several picosecond timescales, making them inadequate for capturing much slower hydration dynamics and its coupling to proton transfer. A common practice in QM/MM simulations of PT processes in proteins such as Complex I, therefore, is to first use standard non-reactive MD runs to capture what appear to be favorable hydrating water structures and then to utilize QM/MM with umbrella sampling over a number of umbrella windows to sample the PT. This approach is to estimate the so-called potential of mean force, or PMF, which is the free energy profile of the excess proton migration along some defined reaction coordinate or “collective variable (CV). However, there are two limitations with this approach. The first is that the water hydration is generally explicitly coupled to the presence of a excess proton in a cooperative fashion and this is largely neglected.^29^ (As will be shown later, our simulations indicate that the rearrangement of water molecules in response to an excess proton can require several hundred picoseconds.) Second, the timescale of the QM/MM sampling in the umbrella windows is generally just a few picoseconds per window given the large computational cost, but the proton hopping between just two waters occurs on that same timescale (∼2 ps in liquid water) and so the QM/MM statistical sampling with such short timescales would be far from complete. (It is not uncommon, e.g., with more efficient reactive MD simulations, to sample in the multiple nanosecond range per umbrella window.^29-35^)

To overcome the limitations of using traditional MD combined with QM/MM as described in the previous paragraph, we have employed multi-state reactive molecular dynamics (MS-RMD)^30^ with explicitly protonatable amino acids. This simulation framework enables explicit modeling of PT via the Grotthuss shuttling mechanism with dynamic protonation of amino acids and has been used extensively to study proton transfer in numerous biomiolecular systems ^29, 31-41^. This approach extends the accessible time and length scales of reactive simulations over QM/MM by 2-3 orders of magnitude, enabling us to capture how complex hydration dynamics are couple to and influence the PT. To quantitatively characterize the hydration, we further applied water wire connectivity order parameter or collective variable (CV), which quantifies the connectivity of water molecules and amino acids along a PT pathway.^29^ Together, these methods provide a powerful platform for probing the interplay between hydration dynamics and proton translocation in Complex I.

In this study, we performed MS-RMD simulations for the Complex I system to investigate proton transfer from the E-channel toward the membrane domain. We focused on the PT pathway spanning E192_ND1_ through D66_ND3_ to E34_ND4L_. To track the location of the excess proton along this path, we applied a curvilinear PT path collective variable (***ξ*** *_*PT*_), and to quantify the hydration environment surrounding the proton, we used a localized water wire connectivity (ϕ *).^29^ The exact definition of both collective variables (CVs) can be found in the Supporting Information. Consistent with previous studies, we found that the segment between D66_ND3_ and E34_ND4L_ exhibits limited hydration without an excess proton. However, our free energy analysis based on the two CVs reveals that PT across this region is not only thermodynamically accessible, but also coupled to the local hydration, highlighting the role of dynamic water networks in regulating proton translocation.

## RESULTS

### Classical MD Simulation and Water Wire Connectivity Analysis

We obtained a stable *Mus musculus* Complex I structure after 1 μs of classical MD simulation. Figure 1A illustrates the equilibrated structure of Complex I embedded in a lipid membrane (the composition and properties of which are given in the Methods section, along with a schematic representation of its overall mechanism. According to the generally accepted mechanism, electrons donated from NADH are transferred through the Fe-S clusters in the peripheral arm, ultimately reducing ubiquinone (Q) to ubiquinol (QH_2_). Subsequently, the reduced quinone migrates toward the entrance of the E-channel, concurrently facilitating proton entry into the E-channel.^6, 7, 9, 11^ To investigate the subsequent proton transfer toward the ND4L subunit, we modeled QH_2_ near the entrance of the E-channel and confirmed that its position is stable during the simulation without explicit restraints. Figure 1B presents the configuration of the QH_2_ and the entrance region of the E-channel from our simulation.

In our simulations, a water network was observed extending from QH_2_, through the E-channel, to the ND4L subunit. Figure 1C shows a representative snapshot of the water network, spanning from E192_ND1_ through D66_ND3_ to E34_ND4L_. This observation is broadly consistent with previous experimental^12, 24-27^ and computational studies;^16, 18^ however, a key advance of this work is the ability to quantitatively assess the hydration of the D66_ND3_–E34_ND4L_ region – previously identified as poorly connected – through thermodynamic analysis of water wire connectivity (ϕ). Note that we use here the water-wire connectivity (ϕ, with no asterisk) that is averaged along the entire path, as these simulations are non-reactive and contain no excess proton. The path for calculating ϕ was constructed from the simulation trajectory by applying a principal curve algorithm^42^ to the positions of water oxygen atoms. The path is represented as 25 equidistant nodes which are shown in Figure 2A. To capture the thermodynamics of the hydration, we ran a set of multiple walkers well-tempered metadynamics simulations^43^ with biasing ϕ. As shown in Figure 2B, the potential of mean force (PMF, i.e., conditional free energy path) of ϕ has a broad well ranging from 0.5 to 0.8. Its minimum is at ϕ = 0.67 (which signifies an incomplete hydration), and there are metastable states ϕ = 0.85 and ϕ = 0.45. Figure 2C shows representative snapshots at two distinct levels of ϕ. Although the ϕ values differ by more than 0.4 between the two states, the overall arrangement of water molecules remains similar, with a few specific water molecules between D66_ND3_ and E34_ND4L_ exhibiting differences. This Indicates that the ϕ is highly sensitive to the arrangement of a few key water molecules in a small region.

**Figure 2.**
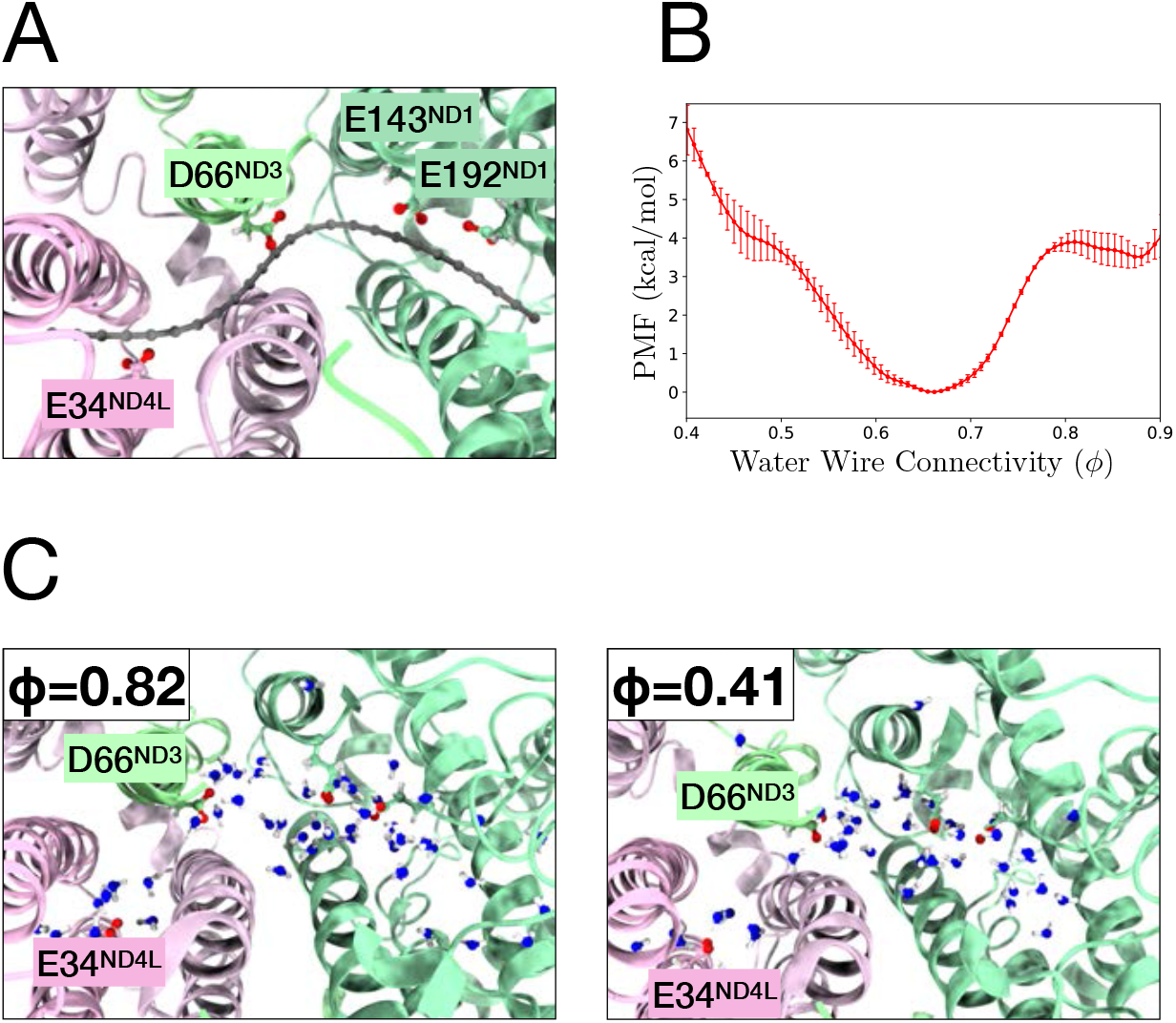
(A) A representative snapshot showing the positions of nodes used to compute water wire connectivity. (B) PMF for water wire connectivity (ϕ) obtained from multiple walkers well-tempered metadynamics simulations. (C) Representative snapshots illustrating the difference in water molecule arrangements at ϕ of 0.82 and 0.41.

Based on these observations, we divided the sampling of the actual PT pathway into two segments and applied umbrella sampling (US) separately for each. For the D66_ND3_–E34_ND4L_ region, which exhibits limited and transient hydration, we performed two-dimensional US using two CVs: the position of the excess proton along the curvilinear PT path (***ξ*** *_*PT*_) and the local water wire connectivity (ϕ *), allowing us to map the free energy surface of the PT reaction. Note that ϕ * is a localized version of water wire connectivity ϕ that quantifies the organization of water molecules near the excess proton by selectively weighting the contribution of nodes near the excess positive charge. In contrast, for the stably hydrated E192_ND1_–D66_ND3_ region, we employed one-dimensional US biasing ***ξ*** *_*PT*_ to compute the PMF as the water connectivity is not an issue in that region. To distinguish these two segments easier, we define ***ξ*** *_*PT*_ by setting the node closest to D66_ND3_ as the origin (***ξ*** *_*PT*_ = 0), with positive values pointing toward E34_ND4L_ and negative values toward E192_ND1_. Although ***ξ*** *_*PT*_ is unitless, we scale it to correspond to distance in Å.

### 2D-Umbrella Sampling MS-RMD simulations for D66-E34

The 2D-PMF of PT in D66_ND3_–E34_ND4L_ region is presented in Figure 3A. The dashed line shows the minimal free energy path (MFEP) of the PT reaction. Note that each umbrella sampling window was sampled for 1–2 ns, and the resulting PMF was averaged over the last four out of eight equally divided blocks. Notably, the PMFs from the first 4 blocks showed significant deviation from the final PMFs, indicating that at least 0.5-1 ns of simulation per US window was required for the convergence of the PMF (Figure S1), which is well beyond the statistical sampling presently accessible by any sort of QM simulation. This result indicates that several hundred picoseconds are needed for water molecules and the excess proton to reorganize within each window and highlights a limitation of QM/MM methods that rely on only a few picoseconds of sampling per umbrella window: This is not enough sampling for the necessary equilibration of water and proton configurations. Even if long classical MD simulations were used to first equilibrate certain water configurations, the presence of an excess proton alters the local hydration thermodynamics.

**Figure 3.**
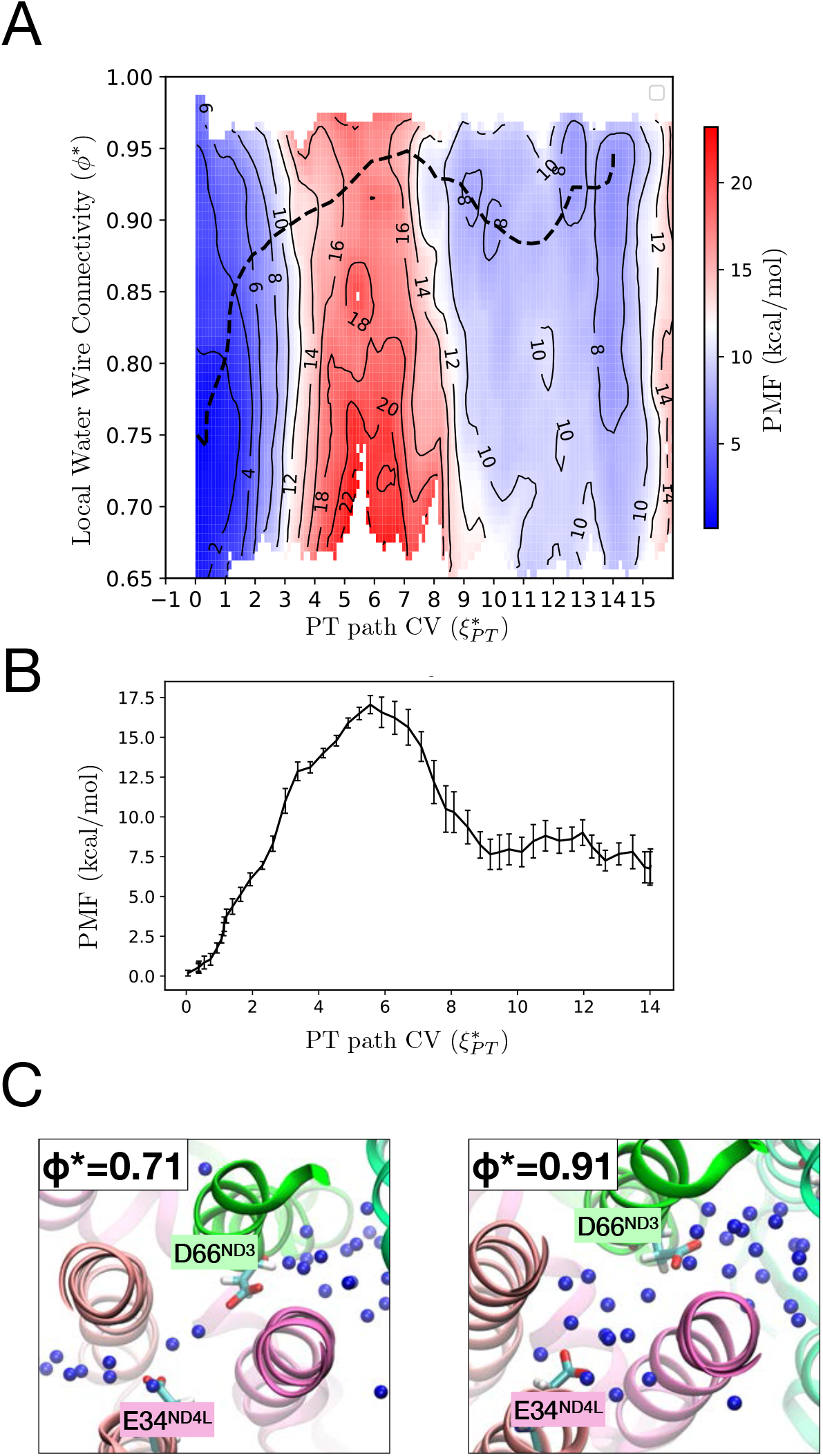
(A) Two-dimensional potential of mean force (2D-PMF) as a function of the local water wire connectivity (ϕ *) and PT path CV (***ξ*** *_*PT*_ ), obtained from 2D umbrella sampling. The minimum free energy path (MFEP) is shown as a dashed line. (B) The PMF of the MFEP projected along the ***ξ*** *_*PT*_. (C) Representative snapshots of water molecule arrangements from two windows at ***ξ*** *_*PT*_ = 5.5, each exhibiting distinct ϕ *, 0.71 and 0.91.

The 2D PMF reveals a coupling between the two CVs in the region near D66_ND3_, where ***ξ*** *_*PT*_ ranges from 0 to 5. When ***ξ*** *_*PT*_ = 0, corresponding to a protonated D66_ND3_, the free energy minimum is at ϕ *≈ 0.75. In contrast, at the saddle point around ***ξ*** *_*PT*_ ≈ 6, ϕ * exceeds 0.9, indicating a fully connected water chain. The MFEP (dashed line) shows that as ***ξ*** *_*PT*_ increases from 0 to 5, ϕ * also increases, indicating explicit coupling between the proton position and hydration^29^ – i.e., as the excess proton moves forward, the less hydrated regions become more easily hydrated. Such proton-induced hydration has been commonly observed in previous studies of other proton channels.^29, 31, 35, 37, 44, 45^ In the context of Complex I, this coupling between proton position and hydration can be a key to regulate proton transfer.

Figure 3B shows the PMF of the MFEP projected on ***ξ*** *_*PT*_, indicating an endergonic proton transfer reaction with an energy barrier (Δ*F*^‡^) of 17.1 ± 0.6 kcal/mol and a reaction free energy (ΔG) of 7 ± 1.0 kcal/mol. Using the classical transition state theory expression, 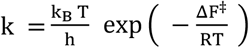, an estimate of the proton transfer rate yields 6.1 s^−1^. Although the this barrier is moderately high, it is comparable to the energy released upon ubiquinone reduction (∼ 800 meV=18.5 kcal/mol). Moreover, as shown in Figure 4A, the E192_ND1_–D66_ND3_ proton transfer is exergonic, further supporting the idea that the overall PT is feasible. In contrast, the 2D PMF allows us to estimate the barrier under low hydrated conditions. When ϕ * = 0.7, the barrier height increases to 22 kcal/mol and the estimated reaction rate is 0.002 s^−1^. This estimated high barrier indicates that proton transfer from D66_ND3_ to E34_ND4L_ is strongly suppressed in less hydrated environments.

**Figure 4.**
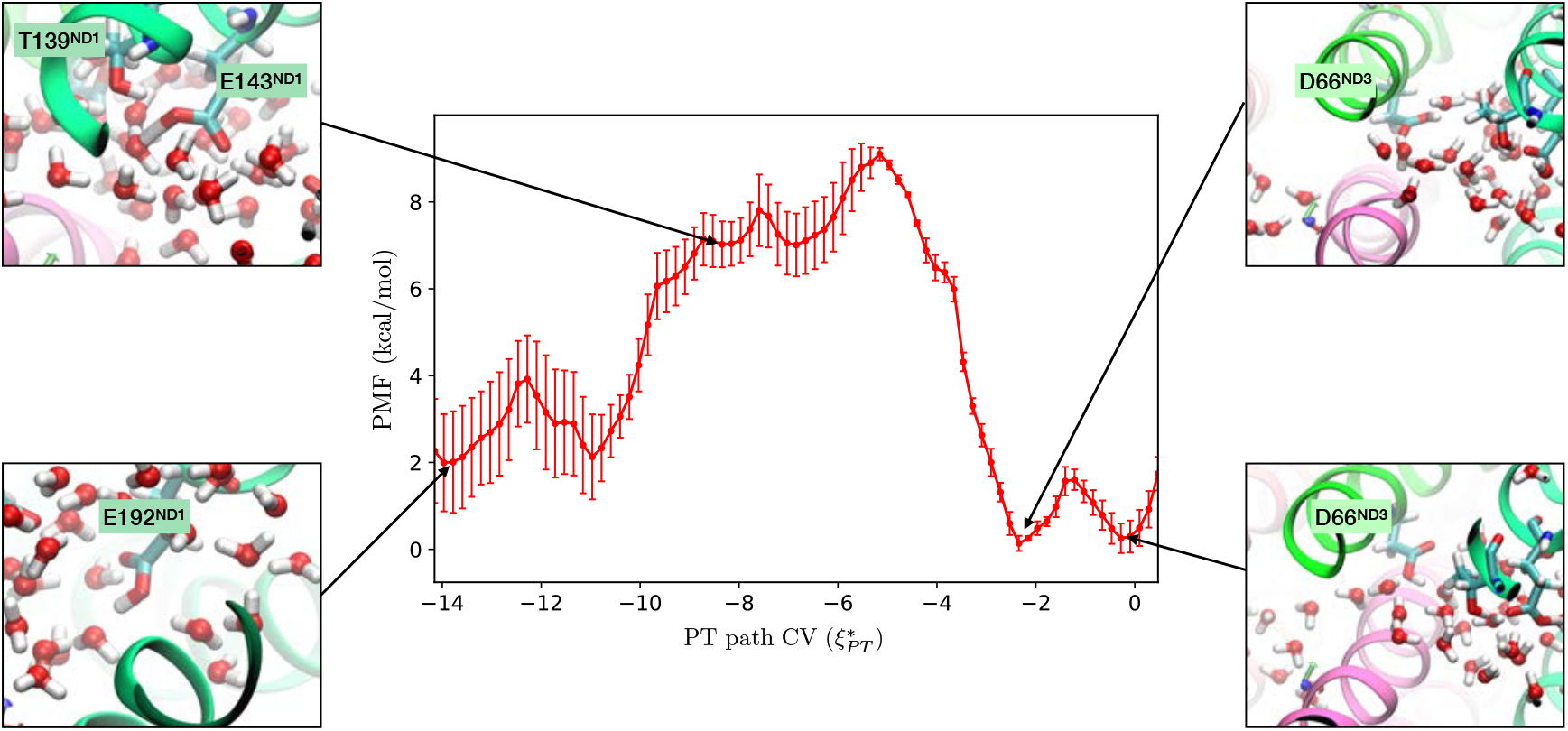
One-dimensional potential of mean force (1D-PMF) for proton transfer along the E192_ND1_–E143_ND1_–D66_ND3_ pathway. Representative snapshots are shown for selected positions along the reaction coordinate, highlighting distinct proton positions and water molecule arrangements.

In a recent paper, Sharma and co-workers reported a QM/MM free-energy barrier of approximately 11 kcal/mol in the same region.^20^ While this similarly identifies a thermodynamic bottleneck, their calculation differs from ours in several important respects. Because of the limited time scale accessible to their QM/MM simulations, the surrounding water molecules remained effectively static, so the energetic cost associated with organizing the water network was not sampled. In addition, their D66_ND3_ side chain was oriented toward E34 at the start of the calculation, whereas our simulations show that D66_ND3_ initially points in the opposite direction. Thus, the rotational rearrangement of this residue – which contributes to the overall barrier in our system – was not captured in their free-energy profile. These methodological differences likely explain the quantitative difference in the barriers while remaining qualitatively consistent.

Given the critical role of hydration in proton transfer, a quantitative descriptor such as water wire connectivity is essential. Figure 3C presents two representative snapshots at ***ξ*** *_*PT*_ ≈ 5.5 with differing ϕ * values. As shown in the left configuration, a few water molecules are aligning between D66_ND3_ and E34_ND4L_ even at less hydrated state (ϕ * = 0.71). In a typical structural assessment, which checks whether water molecules are present within a predefined distance, this configuration can be classified as hydrated state. However, our PMF reveals a significant free energy difference between the two states, and PT is unlikely to proceed in the lower-ϕ * case. By employing ϕ *, we were able to distinguish the thermodynamic disparity between these hydration states.

### 1D-Umbrella Sampling MS-RMD simulation on E192-D66

Compared to the proton transfer from D66_ND3_ to E34_ND4L_, the transfer from E192_ND1_ to D66_ND3_ occurs more readily. As shown in Figure 4, the transfer from E192_ND1_ to D66_ND3_ is exergonic, with a free energy decrease of 2 kcal/mol. The activation barrier (7 kcal/mol) is much lower than 17.1 kcal/mol barrier of transfer from D66_ND3_ to E34_ND4L_, indicating rapid proton transfer to D66_ND3_. The PMF profile along ***ξ*** *_*PT*_ reveals distinct protonation states of residues across the pathway. In the range of ***ξ*** *_*PT*_ = -14 to -10, where E192_ND1_ is protonated, showing two clear local minima corresponding to the protonation of each of carboxylate oxygens. In the region between -2 and 0, D66_ND3_ is protonated as shown in the two right snapshots of Figure 4. In contrast, at ***ξ*** *_*PT*_ = -8 to -6, where the excess charge is near E143_ND1_, there are shallow and less distinct minima indicating that E143_ND1_ is not well protonated, unlike the other acidic residues. The transition state for proton transfer appears between E143_ND1_ and D66_ND3_ at ***ξ*** *_*PT*_ = -5.

A noteworthy finding is that E143_ND1_ is rarely protonated unlike the other Glu and Asp in this PT path. The left top snapshot of Figure 4 shows E143_ND1_ and surrounding water molecules at ***ξ*** *_*PT*_ ≈ -8. A hydronium ion is sufficiently close to protonate E143_ND1_, but its protonated state is not the lowest energy state in the MS-RMD method. Instead, T139_ND1_ donates its hydrogen in a hydrogen bond to E143_ND1_. Due to this hydrogen bond, the pK_a_ of E143_ND1_ decreases; consequently, making it less favorable to be protonated. Importantly, this protonation behavior and its underlying mechanism would not have been accessible through non-reactive MD simulations; it was only revealed by employing MS-RMD, which explicitly accounts for dynamic proton delocalization and transfer events. Interestingly, a recent study did not observe such an interaction between E143_ND1_ and T139_ND1_,^20^ most likely because E143_ND1_ was kept in its neutral form during classical MD simulations. However, E143_ND1_ is expected to be deprotonated during the PT process, and our results suggest that the effect of T139_ND1_ may further lower its pKa, making E143_ND1_ less likely to be in a neutral state than previously assumed.

## DISCUSSION AND CONCLUDING REMARKS

In this work, we have presented a quantitative free energy and kinetic analysis of proton transfer from the E-channel to the ND4L subunit in Complex I. For the first half of the pathway, from E192_ND1_ to D66_ND3_, the proton transfer is exergonic with ΔG = -2 kcal/mol and has barrier of 7 kcal/mol (Figure 4). For the other half from D66_ND3_ to E34_ND4L_, the proton transfer is endergonic with ΔG = 7 kcal/mol and has a significantly higher barrier (Figure 3B). The overall free energy increase for the PT (5 kcal/mol) is smaller compared to the energy release upon the reduction of ubiquinol (18.45 kcal/mol). However, the large free energy barrier between D66_ND3_ and E34_ND4L_ (17 kcal/mol) suggests that the D66_ND3_-E34_ND4L_ region can be a kinetic bottleneck of the PT for the first step of proton pumping in Complex I.

By using the water wire connectivity analysis, we showed that the proton transfer from D66_ND3_ to E34_ND4L_ is highly coupled with the local hydration in the channel. On the other hand, the translocation of an excess proton can stimulate and stabilize the hydration around it, and that higher hydration can then facilitate the PT. Although it is still not clear where the proton entering to the E-channel originates, it has been suggested that two protons can be injected after ubiquinone moves to the entrance of the E-channel.^4^ Our simulations suggest that these protons can promote channel hydration and thereby enable PT that would otherwise be inaccessible.

Our simulations include only a single excess proton, although mechanistic models have suggested that two protons may enter the E-channel during the catalytic cycle. This choice was made intentionally for two key reasons. First, tracking a single proton along the pathway allows clear identification of the coupling between proton position and local hydration, without complex proton–proton interactions. Second, there is currently no evidence that two protons are transferred simultaneously or occupy adjacent positions in the channel. On the contrary, electrostatic repulsion would likely prevent two excess protons from residing in close proximity, even with hydration. As such, we can propose a mechanism based on our PMF results: protonation of D66_ND3_ corresponds to a local minimum on the free energy surface, suggesting that a proton arriving at D66_ND3_ is likely to remain there for an extended time. If a second proton subsequently approaches this region, it would experience an elevated free energy due to the presence of the first proton. In effect, this might reduce the free energy barrier for the second proton to proceed along the pathway. Exploring such cooperative or sequential effects of multiple protons presents an intriguing direction for future research.

While previous studies have suggested possible proton transfer pathways in Complex I,^6, 12, 17, 20, 22^ they have not captured the hydration-dependent behavior of the D66_ND3_–E34_ND4L_ region, which we address here using extended-sampling MS-RMD and water wire connectivity analysis. Although possible transient water network in this region has been reported in some experimental studies,^12, 24-27^ those measurements provide time-averaged water densities and cannot quantify how an excess proton dynamically and transiently reshapes hydration and facilitates transfer. A recent QM/MM study likewise reported proton transfer across the E192–E143–D66–E34 segment, supporting the view that this region constitutes the principal conduit in the E-channel.^20^ Their analysis primarily examined how variations in protonation states modulate the proton-transfer energetics. However, with QM/MM sampling limited to only a few picoseconds per umbrella-sampling window, the water within the channel is virtually immobile on that timescale, making it unlikley to capture the excess proton-induced hydration dynamics. By employing nanosecond timescales and a quantitative analysis of water-wire connectivity, our MS-RMD simulations can capture these dynamics and delineate their thermodynamic consequences. It is also worth noting that recent advances in SCC-DFTB–based QM/MM methods,^46-48^ together with emerging machine-learning strategies for improving semiempirical Hamiltonians,^49, 50^ suggest that future QM/MM approaches may approach longer-timescale sampling that will be needed to serve as a useful complement for studying hydration-coupled PT dynamics in Complex I.

In summary, we have demonstrated that PT through the ND1-ND4L subunits in Complex I is coupled with local hydration, and that the excess proton itself promotes hydration in otherwise dry regions. Our free energy calculations reveal a substantial kinetic barrier in the D66_ND3_–E34_ND4L_ segment, which may serve as a regulatory gate in the overall proton pumping mechanism. These results suggest that hydration is not just a passive environmental factor, but an active component of proton gating and subsequent PT. This work also highlights the importance of treating hydration as a dynamic variable in mechanistic studies of PT in bioenergetic complexes and provides a computational framework that can be extended to other proton transporting proteins.

## METHODS

### Classical MD and MS-RMD Simulation Details

The initial configuration was prepared from the cryo-EM structure of active *Mus musculus* Complex I [PDB ID: 8OM1].^27^ The entire subunits of Complex I were placed in a 1:2:2 of cardiolipin:POPC:POPE membrane and solvated with ∼ 450k TIP3P water molecules. 150 mM NaCl ions were added to neutralize the charge. FMN and Fe-S clusters were placed in the same position of the original structure, and the position of ubiquinone was modeled to be located at the entrance of E-channel, as identified in previous studies.^4, 28, 51-53^ The CHARMM36m (CHARMM36) force field was used for the protein (lipids). For FMN, Fe-S clusters, and ubiquinone, CHARMM style parameters were implemented from Ref. ^54-56^ respectively. To get the equilibrated structure, 1 μs of MD simulation in constant NpT ensemble at 310 K was done with the GROMACS^57^ simulation package with GPU acceleration. The protein backbones were fixed to their original coordinates for the first 300 ns of the simulation to relax the lipid conformation without disrupting protein packing. Subsequently, the position restraint was removed for the remaining 700 ns of the simulation.

After the classical MD simulation, we calculated the water wire connectivity (ϕ), as described in the following section. To ensure sufficient hydration, the configuration with the highest ϕ value within the last 100 ns of the simulation was selected and converted into the MS-RMD initial structure. MS-RMD simulation was done with RAPTOR^30^ module implemented in the LAMMPS^58^ MD package. In MS-RMD simulation, all the TIP3P waters are replaced by MS-EVB 3.2^59^ waters and the selected residues, E202_ND1_, E227_ND1_, E192_ND1_, E143_ND1_, D66_ND3_ and E34_ND4L_, were modeled as EVB-active, using the parameters reported in Ref. ^38, 39^.

To calculate the PMF along ϕ without excess proton, we performed eight walkers well-tempered metadynamics simulation^60, 61^ with LAMMPS and PLUMED simulation packages. Gaussian hills were deposited every 1,000 steps with a height of 0.6 kcal/mol and a width (σ) of 0.02 along the ϕ. The simulations were carried out at 310 K with a bias factor of 35. The walkers shared the biasing potential by reading hill files every 100 steps, allowing enhanced sampling of the free energy landscape. Each walker was simulated for 7.5 ns, resulting in a total simulation time of 60 ns across 8 walkers.

### Collective Variables

In this study, we introduce two CVs to show how the transport of excess protons depend on the hydration of the channel: the water wire connectivity (ϕ and ϕ *) and the curvilinear PT path CV (***ξ*** *_***CEC***_). The definition of both CVs can be found in the Supporting Information. Note that in this study we used two versions of water wire connectivity: the averaged water wire connectivity, ϕ, that represents the geometric mean of the connectivity of all nodes along the path and the local water wire connectivity, ϕ *, that assigns greater weight to the connectivity of nodes near the excess proton.

### Umbrella Sampling and WHAM

Umbrella sampling (US) was performed with the modified version of PLUMED v2.4.^29, 62^ The system was integrated with a 1-fs timestep and the Nose-Hoover thermostat in the constant NVT ensemble at 310 K. For the 2D-US simulations of ***ξ*** *_*PT*_ and ϕ *, a total of 130 US windows were used, with each window running for 1–2 ns, resulting in a total simulation time of over 200 ns. Harmonic force constants of 2500 kcal/mol and 10–30 kcal/mol were applied to ϕ * and ***ξ*** * _*PT*_, respectively. For the 1D-US simulation of the PT path CV (***ξ*** *_*PT*_), 24 windows were used, with each window running for 1-1.5 ns, and force constants ranging from 10–30 kcal/mol were applied. The initial configuration for each window was created by dragging the excess proton along the path. The PMF of PT along the path was computed using the weighted histogram analysis method (WHAM).^63^ To ensure the convergence of the PMF, we performed block averaging for each umbrella sampling window. For the 2D-US calculations, the PMF was averaged over the last four of eight blocks. For the 1D-US calculations, the average was taken over the last three of six blocks.

## Supporting Information

The following files are available free of charge: Supporting figures, detailed descriptions of the simulation and free energy calculation methods, and the mathematical details of the collective variables (PDF).

## Author Contributions

The research was conceived, carried out, and the manuscript was written through contributions of both authors. All authors have given approval to the final version of the manuscript.

## ACKNOWLEDGMENTS

This work was supported by the US Department of Energy, Office of Science, Basic Energy Sciences, under award DE-SC0023318. We acknowledge the use of the Beagle-3 computing cluster funded through the National Institutes of Health by grant 1S10OD028655 (B. Roux, PI), as well as the University of Chicago Research Computing Center (RCC) for computational resources used in this work. This work also used Bridges-2 at the Pittsburgh Supercomputing Center through allocation MCA94P017 and EXPANSE at the San Diego Supercomputer Center through allocation SLC215 from the Advanced Cyberinfrastructure Coordination Ecosystem: Services & Support (ACCESS) program, which is supported by National Science Foundation grants #2138259, #2138286, #2138307, #2137603, and #2138296.

## Table of Contents Graphic

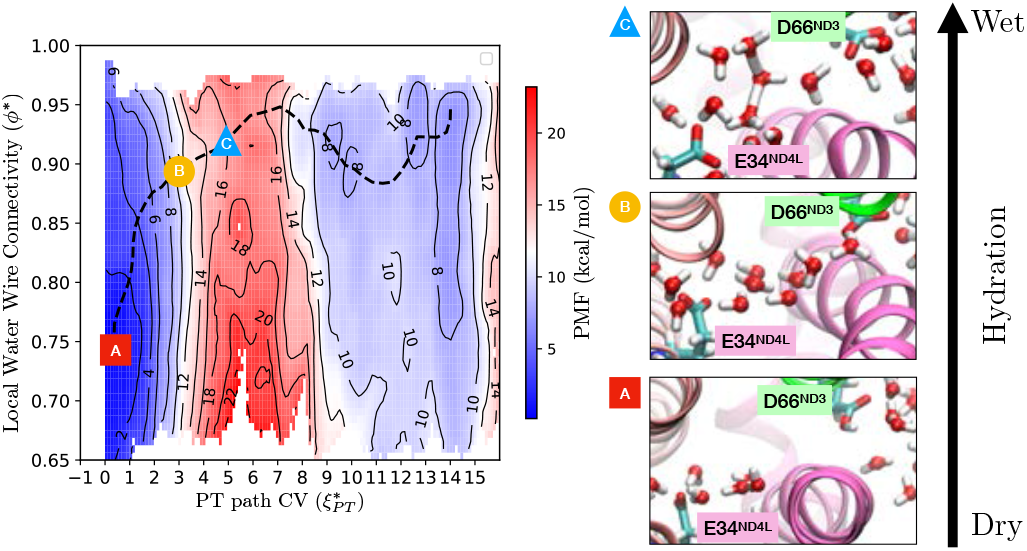

## REFERENCES

(1) Hirst, J. Mitochondrial Complex I. Annual Review of Biochemistry 2013, 82 (1), 551–575. DOI: 10.1146/annurev-biochem-070511-103700

(2) Sazanov, L. A. A giant molecular proton pump: structure and mechanism of respiratory complex I. Nature Reviews Molecular Cell Biology 2015, 16 (6), 375–388.

(3) Brandt, U. Energy Converting NADH: Quinone Oxidoreductase (Complex I). Annual Review of Biochemistry 2006, 75 (1), 69–92.

(4) Djurabekova, A.; Lasham, J.; Zdorevskyi, O.; Zickermann, V.; Sharma, V. Long-range electron proton coupling in respiratory complex I — insight s from molecular simulations of the quinone chamber and antiporter-lik e subunits. Biochemical Journal 2024, 481 (7), 499–514.

(5) Wikström, M.; Sharma, V.; Kaila, V. R. I.; Hosler, J. P.; Hummer, G. New Perspectives on Proton Pumping in Cellular Respiration. Chemical Reviews 2015, 115 (5), 2196–2221.

(6) Sharma, V.; Belevich, G.; Gamiz-Hernandez, A. P.; Róg, T.; Vattulainen, I.; Verkhovskaya, M. L.; Wikström, M.; Hummer, G.; Kaila, V. R. I. Redox-induced activation of the proton pump in the respiratory complex I. Proceedings of the National Academy of Sciences 2015, 112 (37), 11571–11576.

(7) Gamiz-Hernandez, A. P.; Jussupow, A.; Johansson, M. P.; Kaila, V. R. I. Terminal Electron– Proton Transfer Dynamics in the Quinone Reduction of Respiratory Complex I. Journal of the American Chemical Society 2017, 139 (45), 16282–16288.

(8) Kaila, V. R. I. Long-range proton-coupled electron transfer in biological energy conve rsion: towards mechanistic understanding of respiratory complex I. Journal of The Royal Society Interface 2018, 15 (141), 20170916.

(9) Gu, J.; Liu, T.; Guo, R.; Zhang, L.; Yang, M. The coupling mechanism of mammalian mitochondrial complex I. Nature Structural & Molecular Biology 2022, 29 (2), 172–182.

(10) Mühlbauer, M. E.; Saura, P.; Nuber, F.; Di Luca, A.; Friedrich, T.; Kaila, V. R. I. Water-Gated Proton Transfer Dynamics in Respiratory Complex I. Journal of the American Chemical Society 2020, 142 (32), 13718–13728.

(11) Bridges, H. R.; Fedor, J. G.; Blaza, J. N.; Di Luca, A.; Jussupow, A.; Jarman, O. D.; Wright, J. J.; Agip, A.-N. A.; Gamiz-Hernandez, A. P.; Roessler, M. M.; et al. Structure of inhibitor-bound mammalian complex I. Nature Communications 2020, 11 (1).

(12) Parey, K.; Lasham, J.; Mills, D. J.; Djurabekova, A.; Haapanen, O.; Yoga, E. G.; Xie, H.; Kühlbrandt, W.; Sharma, V.; Vonck, J.; et al. High-resolution structure and dynamics of mitochondrial complex I—Insi ghts into the proton pumping mechanism. Science Advances 2021, 7 (46), eabj3221.

(13) Agip, A.-N. A.; Blaza, J. N.; Bridges, H. R.; Viscomi, C.; Rawson, S.; Muench, S. P.; Yang, J. Cryo-EM structures of complex I from mouse heart mitochondria in two b iochemically defined states. Nature Structural & Molecular Biology 2018, 25 (7), 548–556.

(14) Baradaran, R.; Berrisford, J. M.; Minhas, G. S.; Sazanov, L. A. Crystal structure of the entire respiratory complex I. Nature 2013, 494 (7438), 443–448.

(15) Chung, I.; Serreli, R.; Cross, J. B.; Di Francesco, M. E.; Marszalek, J. R.; Hirst, J. Cork-in-bottle mechanism of inhibitor binding to mammalian complex I. Science Advances 2021, 7 (20), eabg4000.

(16) Kim, H.; Saura, P.; Pöverlein, M. C.; Gamiz-Hernandez, A. P.; Kaila, V. R. I. Quinone Catalysis Modulates Proton Transfer Reactions in the Membrane Domain of Respiratory Complex I. Journal of the American Chemical Society 2023, 145 (31), 17075–17086.

(17) Röpke, M.; Saura, P.; Riepl, D.; Pöverlein, M. C.; Kaila, V. R. I. Functional Water Wires Catalyze Long-Range Proton Pumping in the Mamma lian Respiratory Complex I. Journal of the American Chemical Society 2020, 142 (52), 21758–21766.

(18) Röpke, M.; Riepl, D.; Saura, P.; Di Luca, A.; Mühlbauer, M. E.; Jussupow, A.; Gamiz-Hernandez, A. P.; Kaila, V. R. I. Deactivation blocks proton pathways in the mitochondrial complex I. Proceedings of the National Academy of Sciences 2021, 118 (29), e2019498118.

(19) Di Luca, A.; Gamiz-Hernandez, A. P.; Kaila, V. R. I. Symmetry-related proton transfer pathways in respiratory complex I. Proceedings of the National Academy of Sciences 2017, 114 (31).

(20) Simsive, L.; Zdorevskyi, O.; Sharma, V. Proton Transfer through a Charged Conduit in Respiratory Complex I: Long-Range Effects and Conformational Gating. Journal of Chemical Information and Modeling 2025, 65 (19), 10600–10612.

(21) Agip, A.-N. A.; Blaza, J. N.; Fedor, J. G.; Hirst, J. Mammalian Respiratory Complex I Through the Lens of Cryo-EM. Annual Review of Biophysics 2019, 48 (1), 165–184.

(22) Kravchuk, V.; Petrova, O.; Kampjut, D.; Wojciechowska-Bason, A.; Breese, Z.; Sazanov, L. A universal coupling mechanism of respiratory complex I. Nature 2022, 609 (7928), 808–814.

(23) Zdorevskyi, O.; Djurabekova, A.; Lasham, J.; Sharma, V. Horizontal proton transfer across the antiporter-like subunits in mito chondrial respiratory complex I. Chemical Science 2023, 14 (23), 6309–6318.

(24) Grba, D. N.; Hirst, J. Mitochondrial complex I structure reveals ordered water molecules for catalysis and proton translocation. Nature Structural & Molecular Biology 2020, 27 (10), 892–900.

(25) Wang, P.; Demaray, J.; Moroz, S.; Stuchebrukhov, A. A. Searching for proton transfer channels in respiratory complex I. Biophysical Journal 2024, S0006349524005186.

(26) Chung, I.; Wright, J. J.; Bridges, H. R.; Ivanov, B. S.; Biner, O.; Pereira, C. S.; Arantes, G. M.; Hirst, J. Cryo-EM structures define ubiquinone-10 binding to mitochondrial compl ex I and conformational transitions accompanying Q-site occupancy. Nature Communications 2022, 13 (1), 2758.

(27) Grba, D. N.; Chung, I.; Bridges, H. R.; Agip, A.-N. A.; Hirst, J. Investigation of hydrated channels and proton pathways in a high-resol ution cryo-EM structure of mammalian complex I. Science Advances 2023, 9 (31), eadi1359.

(28) Haapanen, O.; Sharma, V. Role of water and protein dynamics in proton pumping by respiratory co mplex I. Scientific Reports 2017, 7 (1), 7747.

(29) Li, C.; Voth, G. A. A quantitative paradigm for water-assisted proton transport through proteins and other confined spaces. Proceedings of the National Academy of Sciences of the United States o f America 2021, 118 (49), 1–8.

(30) Kaiser, S.; Yue, Z.; Peng, Y.; Nguyen, T. D.; Chen, S.; Teng, D.; Voth, G. A. Molecular Dynamics Simulation of Complex Reactivity with the Rapid App roach for Proton Transport and Other Reactions (RAPTOR) Software Packa ge. The Journal of Physical Chemistry B 2024, 128 (20), 4959–4974.

(31) Li, C.; Yue, Z.; Espinoza-Fonseca, L. M.; Voth, G. A. Multiscale Simulation Reveals Passive Proton Transport Through SERCA on the Microsecond Timescale. Biophysical Journal 2020, 119 (5), 1033–1040.

(32) Liu, Y.; Li, C.; Gupta, M.; Verma, N.; Johri, A. K.; Stroud, R. M.; Voth, G. A. Key computational findings reveal proton transfer as driving the functional cycle in the phosphate transporter PiPT. Proceedings of the National Academy of Sciences 2021, 118 (25), e2101932118.

(33) Liu, Y.; Li, C.; Gupta, M.; Stroud, R. M.; Voth, G. A. Kinetic network modeling with molecular simulation inputs: A proton-coupled phosphate symporter. Biophysical Journal 2024, 123 (24), 4191–4199.

(34) Liang, R.; Swanson, J. M. J.; Madsen, J. J.; Hong, M.; DeGrado, W. F.; Voth, G. A. Acid activation mechanism of the influenza A M2 proton channel. Proceedings of the National Academy of Sciences 2016, 113 (45), E6955–E6964.

(35) Watkins, L. C.; DeGrado, W. F.; Voth, G. A. Multiscale Simulation of an Influenza A M2 Channel Mutant Reveals Key Features of Its Markedly Different Proton Transport Behavior. Journal of the American Chemical Society 2022, 144 (2), 769–776.

(36) Lee, S.; Mayes, H. B.; Swanson, J. M. J.; Voth, G. A. The Origin of Coupled Chloride and Proton Transport in a Cl–/H+ Antipo rter. Journal of the American Chemical Society 2016, 138 (45), 14923–14930.

(37) Lee, S.; Swanson Jessica, M. J.; Voth Gregory, A. Multiscale Simulations Reveal Key Aspects of the Proton Transport Mech anism in the ClC-ec1 Antiporter. Biophysical Journal 2016, 110 (6), 1334–1345.

(38) Li, C.; Voth, G. A. Accurate and Transferable Reactive Molecular Dynamics Models from Cons trained Density Functional Theory. The Journal of Physical Chemistry B 2021, 125 (37), 10471–10480.

(39) Zuchniarz, J.; Liu, Y.; Li, C.; Voth, G. A. Accurate pKa Calculations in Proteins with Reactive Molecular Dynamics Provide Physical Insight Into the Electrostatic Origins of Their Values. The Journal of Physical Chemistry B 2022, 126 (38), 7321–7330.

(40) Liu, Y.; Li, C.; Voth, G. A. Generalized Transition State Theory Treatment of Water-Assisted Proton Transport Processes in Proteins. The Journal of Physical Chemistry B 2022, 126 (49), 10452–10459.

(41) Liu, Y.; Li, C.; Freites, J. A.; Tobias, D. J.; Voth, G. A. Quantitative insights into the mechanism of proton conduction and selectivity for the human voltage-gated proton channel Hv1. Proceedings of the National Academy of Sciences 2024, 121 (38), e2407479121.

(42) Hastie, T.; Stuetzle, W. Principal Curves. Journal of the American Statistical Association 1989, 84 (406), 502–516.

(43) Raiteri, P.; Laio, A.; Gervasio, F. L.; Micheletti, C.; Parrinello, M. Efficient Reconstruction of Complex Free Energy Landscapes by Multiple Walkers Metadynamics. The Journal of Physical Chemistry B 2006, 110 (8), 3533–3539.

(44) Li, C.; Yue, Z.; Newstead, S.; Voth, G. A. Proton coupling and the multiscale kinetic mechanism of a peptide tran sporter. Biophysical Journal 2022, 121 (12), 2266–2278.

(45) Liang, R.; Swanson, J. M. J.; Peng, Y.; Wikström, M.; Voth, G. A. Multiscale simulations reveal key features of the proton-pumping mechanism in cytochrome c oxidase. Proceedings of the National Academy of Sciences 2016, 113 (27), 7420–7425.

(46) Cui, Q.; Elstner, M.; Kaxiras, E.; Frauenheim, T.; Karplus, M. A QM/MM Implementation of the Self-Consistent Charge Density Functional Tight Binding (SCC-DFTB) Method. The Journal of Physical Chemistry B 2001, 105 (2), 569–585.

(47) Gaus, M.; Cui, Q.; Elstner, M. DFTB3: Extension of the Self-Consistent-Charge Density-Functional Tight-Binding Method (SCC-DFTB). Journal of Chemical Theory and Computation 2011, 7 (4), 931–948.

(48) Kubař, T.; Elstner, M.; Cui, Q. Hybrid quantum mechanical/molecular mechanical methods for studying energy transduction in biomolecular machines. Annual review of biophysics 2023, 52 (1), 525–551.

(49) Giese, T. J.; Zeng, J.; Lerew, L.; McCarthy, E.; Tao, Y.; Ekesan, Ş.; York, D. M. Software Infrastructure for Next-Generation QM/MM−ΔMLP Force Fields. The Journal of Physical Chemistry B 2024, 128 (26), 6257–6271.

(50) Zeng, J.; Giese, T. J.; Ekesan, Ş.; York, D. M. Development of Range-Corrected Deep Learning Potentials for Fast, Accurate Quantum Mechanical/Molecular Mechanical Simulations of Chemical Reactions in Solution. Journal of Chemical Theory and Computation 2021, 17 (11), 6993–7009.

(51) Warnau, J.; Sharma, V.; Gamiz-Hernandez, A. P.; Di Luca, A.; Haapanen, O.; Vattulainen, I.; Wikström, M.; Hummer, G.; Kaila, V. R. I. Redox-coupled quinone dynamics in the respiratory complex I. Proceedings of the National Academy of Sciences 2018, 115 (36), E8413–E8420.

(52) Hoias Teixeira, M.; Menegon Arantes, G. Balanced internal hydration discriminates substrate binding to respiratory complex I. Biochimica et Biophysica Acta (BBA) - Bioenergetics 2019, 1860 (7), 541–548.

(53) Gupta, C.; Khaniya, U.; Chan, C. K.; Dehez, F.; Shekhar, M.; Gunner, M. R.; Sazanov, L.; Chipot, C.; Singharoy, A. Charge Transfer and Chemo-Mechanical Coupling in Respiratory Complex I. Journal of the American Chemical Society 2020, 142 (20), 9220–9230.

(54) Galkin, A.; Dröse, S.; Brandt, U. The proton pumping stoichiometry of purified mitochondrial complex I r econstituted into proteoliposomes. Biochimica et Biophysica Acta (BBA) - Bioenergetics 2006, 1757 (12), 1575–1581.

(55) Chang, C. H.; Kim, K. Density Functional Theory Calculation of Bonding and Charge Parameters for Molecular Dynamics Studies on [FeFe] Hydrogenases. Journal of Chemical Theory and Computation 2009, 5 (4), 1137–1145.

(56) Kim, S.; Lee, J.; Jo, S.; Brooks, C. L.; Lee, H. S.; Im, W. CHARMM-GUI ligand reader and modeler for CHARMM force field generation of small molecules. Journal of Computational Chemistry 2017, 38 (21), 1879–1886.

(57) Abraham, M. J.; Murtola, T.; Schulz, R.; Páll, S.; Smith, J. C.; Hess, B.; Lindahl, E. GROMACS: High performance molecular simulations through multi-level pa rallelism from laptops to supercomputers. SoftwareX 2015, 1-2, 19–25.

(58) Thompson, A. P.; Aktulga, H. M.; Berger, R.; Bolintineanu, D. S.; Brown, W. M.; Crozier, P. S.; In ‘T Veld, P. J.; Kohlmeyer, A.; Moore, S. G.; Nguyen, T. D.; et al. LAMMPS - a flexible simulation tool for particle-based materials model ing at the atomic, meso, and continuum scales. Computer Physics Communications 2022, 271, 108171.

(59) Biswas, R.; Tse, Y.-L. S.; Tokmakoff, A.; Voth, G. A. Role of Presolvation and Anharmonicity in Aqueous Phase Hydrated Proto n Solvation and Transport. The Journal of Physical Chemistry B 2016, 120 (8), 1793–1804.

(60) Dama, J. F.; Parrinello, M.; Voth, G. A. Well-Tempered Metadynamics Converges Asymptotically. Physical Review Letters 2014, 112 (24), 240602.

(61) Barducci, A.; Bussi, G.; Parrinello, M. Well-Tempered Metadynamics: A Smoothly Converging and Tunable Free-Energy Method. Physical Review Letters 2008, 100 (2), 020603.

(62) Tribello, G. A.; Bonomi, M.; Branduardi, D.; Camilloni, C.; Bussi, G. PLUMED 2: New feathers for an old bird. Computer Physics Communications 2014, 185 (2), 604–613.

(63) Kumar, S.; Rosenberg, J. M.; Bouzida, D.; Swendsen, R. H.; Kollman, P. A. THE weighted histogram analysis method for free-energy calculations on biomolecules. I. The method. Journal of Computational Chemistry 1992, 13 (8), 1011–1021.

