## Supplementary Information for "Hydration-Controlled Proton Transport in Respiratory Complex I"

### <u>for</u>

### <u>Supporting Figures</u>

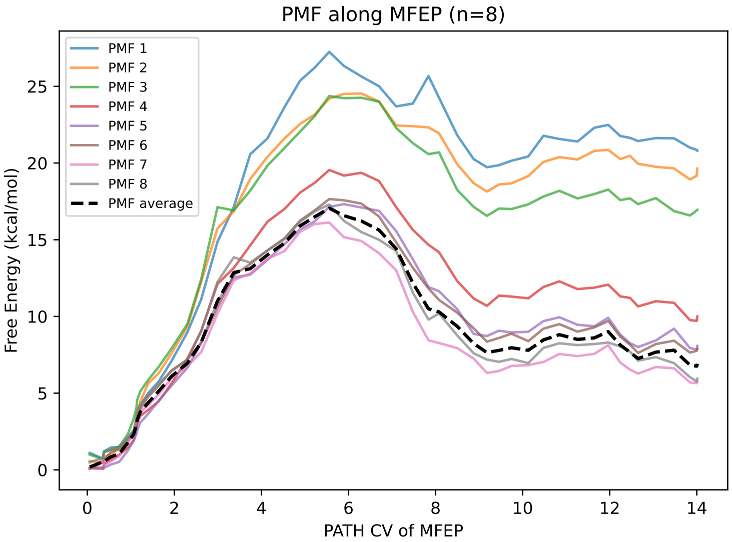


**Figure S1** The PMF results for each of the eight blocks along the MFEP are shown, with the average of the last four blocks (corresponding to Figure 3B) indicated as a dashed line.

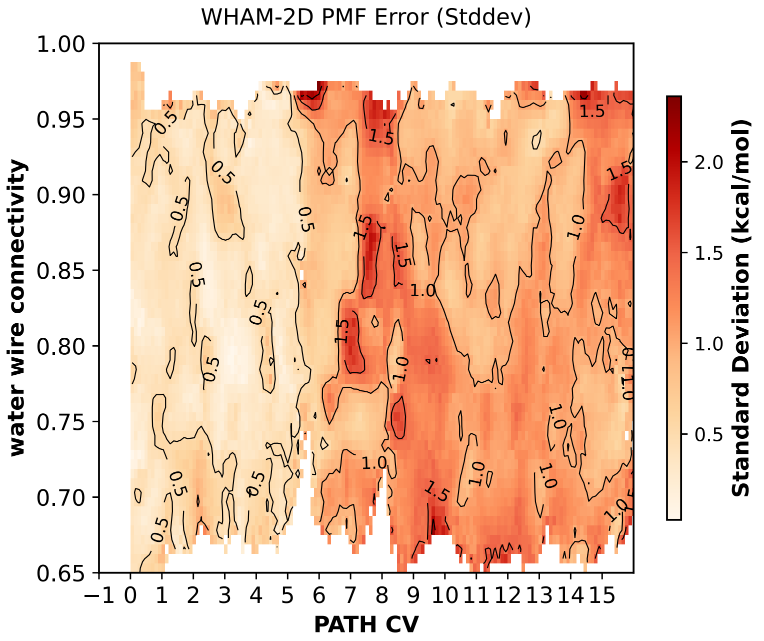


**Figure S2** The standard deviation of the 2D-PMF over the last four blocks

### MS-RMD Simulation and Free Energy Calculation

**Classical MD.** The initial configuration of the system was prepared from the cryo-EM structure of the active *Mus musculus* Complex I (PDB ID: 8OM1).[1] The complete Complex I assembly was embedded in a mixed lipid bilayer composed of cardiolipin, POPC, and POPE in a 1:2:2 molar ratio. The system was solvated with approximately 450,000 TIP3P water molecules, and $Na^+$ and $Cl^-$ ions were added to neutralize the system and achieve a physiological salt concentration of 150 mM. Flavin mononucleotide (FMN) and all iron–sulfur (Fe–S) clusters were retained at their positions in the original cryo-EM structure. The ubiquinone molecule was modeled at the entrance of the E-channel. The CHARMM36m force field was employed for the protein, while lipids were described using the CHARMM36 lipid force field. Parameters for FMN, Fe–S clusters, and ubiquinone were adopted from previously published CHARMM-compatible parameter sets.[2-4]

Classical molecular dynamics simulations were performed using the GROMACS[5] simulation package with GPU acceleration in the NpT ensemble at 310 K. A 2 fs integration time step was used with LINCS algorithm.[6] A Nosé–Hoover thermostat and a Parrinello–Rahman barostat were used. Long-range electrostatics were treated using the particle mesh Ewald (PME) method with a real-space cutoff of 1.2 nm with a Lennard–Jones force-switch from 1.0 to 1.2 nm. To allow relaxation of the membrane environment without perturbing the protein packing, positional restraints were applied to the protein backbone atoms for the first 300 ns of the equilibration. These restraints were subsequently removed, and the system was further equilibrated for an additional 700 ns, yielding a total equilibration time of 1 μs. The equilibrated structure obtained at the end of this protocol was used as the starting point for subsequent MS-RMD simulations.

**MS-RMD.** In the MS-RMD methodology,[7] the Hamiltonian of the system is defined with respect to a basis of diabatic states, or bonding topologies, as

**Commented [GV1]:** Cite Scott's paper here

$$\mathbf{H}^{\mathrm{RMD}} = \boldsymbol{\Sigma}_{ij} \, |i\rangle h_{ij} \langle j| \tag{S1}$$

where $|i\rangle$ represents diabatic state $i$, $h_{ii}$ is the classical force field potential energy of $|i\rangle$, and $h_{ij}$ is a coupling term corresponding to the transition probability between the two states, all dependent on a particular set of nuclear coordinates. The ground state energy of the system is then the lowest eigenvalue of

$$\mathbf{H}^{\mathrm{RMD}}\mathbf{c} = \mathrm{E}\mathbf{c} \tag{S2}$$

where each eigenvector $\mathbf{c}$ has components $c_i$ representing the contribution of each diabatic state to the linear combination, and the ground state is the eigenvector associated with that eigenvalue.

**Umbrella Sampling and Free Energy Reconstruction**. Umbrella sampling (US) simulations were performed using a modified version of PLUMED v2.4[8] interfaced with the MS-RMD framework. The equations of motion were integrated with a 1 fs time step in the NVT ensemble at 310 K, with temperature controlled using a Nosé–Hoover thermostat.

For the two-dimensional umbrella sampling (2D-US) calculations along the proton transport path collective variable, $\boldsymbol{\xi}*_{PT}$, and the localized water wire connectivity, $\phi *$, umbrella window centers were spaced by 0.17–0.33 along $\boldsymbol{\xi}*_{PT}$, and 0.05 along $\phi *$, and a total of 170 umbrella windows were employed. Each window was simulated for 1–2 ns, resulting in an aggregate sampling time exceeding 200 ns. Harmonic biasing potentials were applied independently to each collective variable, with force constants of 2500 kcal/mol for $\phi *$ and 10–30 kcal/mol for $\boldsymbol{\xi}*_{PT}$. The window centers were distributed to ensure sufficient overlap in both collective variables across the sampled region.

For the one-dimensional umbrella sampling (1D-US) calculations along the proton transport path coordinate $\boldsymbol{\xi}*_{PT}$, 24 umbrella windows were used. Each window was simulated for 1–1.5 ns with harmonic force constants ranging from 10 to 30 kcal/mol. Initial configurations for each umbrella window were generated by gradually dragging the excess proton along the predefined curvilinear path to the target window center.

Free energy profiles were reconstructed using the weighted histogram analysis method (WHAM). Convergence of the PMFs was assessed using block averaging. For the 2D-US simulations, the trajectories were divided into eight equal time blocks, and the PMF was averaged over the final four blocks. For the 1D-US simulations, the PMF was averaged over the final three of six blocks. Statistical uncertainty was estimated from the standard deviation across the converged blocks, as reported in the Supporting Figures.

**Convergence of 2D US PMFs.** To confirm the convergence of 2D PMFs, we calculated PMFs for the eight equally divided blocks. Figure S1 presents a comparison of the eight 2D PMFs projected along the minimum free energy path (MFEP). PMF 1 through PMF 8 correspond to results obtained from eight time-ordered blocks of the simulations. The dashed line represents the average PMF calculated from the last four blocks. Notably, the PMFs from the first three blocks exhibit significantly higher energy compared to the later blocks, likely because the water molecules in each window had not yet reached an optimal configuration. Although the fourth block appears closer to convergence, it still shows elevated energy. Therefore, only the final four blocks were considered converged and used to compute the average. Since each umbrella sampling window underwent 1–2 ns of simulation, the final four blocks represent data collected after 0.5–1 ns, indicating that the convergence occurs on this timescale.

The standard deviation across the last four 2D PMF blocks is shown in Figure S2. Within the region of MFEP, the deviation remains within approximately 1 kcal/mol, indicating satisfactory convergence of the free energy surface in this critical region.

**Minimum Free Energy Path**. To extract a smooth and physical minimum free energy path (MFEP) from the two-dimensional PMF F(x,y), we employed a customized nudged-elastic-band (NEB) protocol.[9] The raw surface was first interpolated onto a 300 × 300 grid, and the map was Gaussian-smoothed ($\sigma = 1$ grid unit). Because the energetic span along $\boldsymbol{\xi} *_{PT}$ exceeded that of $\phi *$, the energy gradient was re-weighted with an anisotropic factor ($\lambda_{aniso} = 2.0$) to avoid an x-axis biased path. Each internal bead then experienced four contributions at every iteration: (i) the rescaled free-energy force, (ii) a spring force that equalized inter-bead spacing ($k_{spring}$ = 15 kcal/mol), (iii) a modest length-penalty term that shortened the path ($\lambda_{len}$ = 0.18), and (iv) a soft harmonic wall that prevented excursions beyond the PMF grid. Beads were updated by steepest descent with a step size α = 0.004 for 1200 iterations, and the chain was re-parameterized to uniform arclength every ten steps. The termini were fixed at the PMF minima of $\boldsymbol{\xi} *_{PT} \approx 0.0$ and $\boldsymbol{\xi} *_{PT} \approx 14.3$. Convergence was reached when the force dropped below 0.02 kcal/mol.

**Water Wire Connectivity Metadynamics.** To construct the PMF along the water wire connectivity, well-tempered metadynamics in a multiple-walker framework was used. The simulation protocol and parameters were identical to those used in the MS-RMD simulations, except that the RAPTOR reactive framework was disabled and the simulations were carried out in a non-reactive classical MD mode. The water wire connectivity collective variable, ϕ, was biased, with Gaussian bias potentials deposited every 1000 steps using an initial height of 0.6 and a width of 0.02 at 310 K. A bias factor of 35 was employed to ensure smooth convergence. To accelerate convergence, eight walkers were run in parallel, periodically exchanging bias information, allowing efficient cooperative sampling of water wire formation and disruption.

### The Definition of Collective Variables

**Water wire connectivity.** The water wire connectivity, ϕ, is a quantitative measure of how continuously water molecules are connected along a defined path, first introduced in Ref. [10]. It is calculated through multiple steps. First, a principal curve representing the average water wire is discretized into evenly spaced nodes. For each node, the water coordination number is computed as $s_i = \sum_{I=1}^{N_w} f_{sw}(|X_I - X_i|)$. Here, $\{X_i\}$ and $\{X_I\}$ represent the coordination of the nodes and oxygens of water respectively, and $f_{\mathrm{sw}}$ is a smooth, differentiable switching function. This coordination number is then transformed into a smooth occupancy value using a Fermi-type function

Commented [GV2]: Equations should be numbered here (S1), (S2) etc

$$I_i = \frac{1}{1 + \exp\left(-\frac{s_i - s_w}{\sigma}\right)} \quad \text{(S3)}$$

where parameters $s_w$ and σ are constant that indicate the degree to which a node along the path is occupied given its water coordination number.

Next, the pairwise connectivity between adjacent nodes is defined as the average of their occupancy values ($\mathrm{f_{i,i+1}} = (\mathrm{I_i} + \mathrm{I_{i+1}})/2$), and the overall connectivity is evaluated as the geometric mean over all node pairs along the curve.

$$\phi = \left(\prod_{i=1}^{N-1} \mathrm{f_{i,i+1}}\right)^{\frac{1}{N-1}} \quad \text{(S4)}$$

Since ϕ is computed as the geometric mean over all node pairs along the entire path, it can be significantly affected by poor connectivity in even a small segment of the path. However, for proton transport analysis, only the water surrounding the excess proton is particularly critical. To account for this, a localized version of the connectivity, ϕ ∗, is used, as defined

$$\phi * = \left(\prod_{i=1}^{N-1} f_{i,i+1}\right)^{\frac{1}{N-1}}, \quad \text{(S5)}$$

$$\text{where } f_{i,i+1} = \frac{I_i^{a_i} + I_{i+1}^{a_{i+1}}}{2} \text{ and } a_i = f_{\mathrm{SC}}(|\boldsymbol{r}_{\mathrm{CEC}} - X_i|)$$

This expression selectively weights the contribution of each node based on their proximity to the center of excess charge ($\boldsymbol{r}_{\text{CEC}}$), effectively focusing on the region most relevant to proton translocation. The definition of the center of excess charge is in the following section.

**Curvilinear path collective variable.** To define the curvilinear PT path CV, $\boldsymbol{\xi}_{PT}$, it is necessary to determine the position of the center of excess charge (CEC) first; the CEC can be conveniently defined with respect to these MS-RMD quantities as

$$\mathbf{r}_{\text{CEC}} = \sum_i c_i^2 \, \mathbf{r}_i^{\text{COC}} \tag{S6}$$

where $\mathbf{r}_i^{\text{COC}}$ is the center of charge of the protonated species in state $|i\rangle$. To bias the position of CEC on the curvilinear PT path, we used geometrical path function based on Ref. [11].

$$\xi_{PT'} = i_2 + sign(i_2 - i_1) \cdot \frac{\sqrt{(\boldsymbol{v_1} \cdot \boldsymbol{v}_3)^2 - |\boldsymbol{v}_3|^2(|\boldsymbol{v_1}|^2 - |\boldsymbol{v}_2|^2)}}{2|\boldsymbol{v_3}|^2} - \frac{\boldsymbol{v_1} \cdot \boldsymbol{v_3} - |\boldsymbol{v_3}|^2}{2|\boldsymbol{v_3}|^2} \tag{S7}$$

Here, $\mathbf{v}_1$ and $\mathbf{v}_2$ are the vectors connecting from the $\mathbf{r}_{\text{CEC}}$ to the closest and the second closest nodes of the path, respectively, while $i_1$ and $i_2$ are the indices of the nodes. $\mathbf{v}_3$ denotes the vector connecting from the closest node to the second closest node. As it is defined, $\xi_{PT}$ is an unitless variable that depends on the number of nodes and distances between them. We used the same nodes that is used for ϕ, where the distance between nodes is 1.5 Å. For easier comparison, we defined a shifted and rescaled path CV, $\boldsymbol{\xi}*_{PT}$, where the node closest to $\text{D66}_{\text{ND3}}$ was set to $0^{\text{th}}$ node. The direction toward $\text{E34}_{\text{ND4L}}$ was defined as positive, and the direction toward $\text{E192}_{\text{ND1}}$ as negative. The CV was rescaled by a factor of 1.5 to match the unit of Å.

**Data Availability**

Representative initial and final structures of the equilibrated membrane–protein system are provided as GROMACS coordinate files (.gro). For the MS-RMD simulations, representative initial and final configurations are provided in LAMMPS trajectory format. In addition, input files required to reproduce the classical MD equilibration and umbrella sampling calculations are provided. All of this information is publicly available at:

https://github.com/jongho1994/hydration-controlled-proton-transport

**Supplemental References**

(1) Grba, D. N.; Chung, I.; Bridges, H. R.; Agip, A.-N. A.; Hirst, J. Investigation of hydrated channels and proton pathways in a high-resolution cryo-EM structure of mammalian complex I. *Science Advances* **2023**, *9* (31), eadi1359.
(2) Galkin, A.; Dröse, S.; Brandt, U. The proton pumping stoichiometry of purified mitochondrial complex I reconstituted into proteoliposomes. *Biochimica et Biophysica Acta (BBA) - Bioenergetics* **2006**, *1757* (12), 1575-1581.
(3) Chang, C. H.; Kim, K. Density Functional Theory Calculation of Bonding and Charge Parameters for Molecular Dynamics Studies on [FeFe] Hydrogenases. *Journal of Chemical Theory and Computation* **2009**, *5* (4), 1137-1145.
(4) Kim, S.; Lee, J.; Jo, S.; Brooks, C. L.; Lee, H. S.; Im, W. CHARMM-GUI ligand reader and modeler for CHARMM force field generation of small molecules. *Journal of Computational Chemistry* **2017**, *38* (21), 1879-1886.
(5) Abraham, M. J.; Murtola, T.; Schulz, R.; Páll, S.; Smith, J. C.; Hess, B.; Lindahl, E. GROMACS: High performance molecular simulations through multi-level parallelism from laptops to supercomputers. *SoftwareX* **2015**, *1-2*, 19-25.
(6) Hess, B.; Bekker, H.; Berendsen, H. J. C.; Fraaije, J. G. E. M. LINCS: A linear constraint solver for molecular simulations. *Journal of computational chemistry* **1997**, *18* (12), 1463-1472.
(7) Kaiser, S.; Yue, Z.; Peng, Y.; Nguyen, T. D.; Chen, S.; Teng, D.; Voth, G. A. Molecular Dynamics Simulation of Complex Reactivity with the Rapid App roach for Proton Transport and Other Reactions (RAPTOR) Software Packa ge. *The Journal of Physical Chemistry B* **2024**, *128* (20), 4959-4974
(8) Tribello, G. A.; Bonomi, M.; Branduardi, D.; Camilloni, C.; Bussi, G. PLUMED 2: New feathers for an old bird. *Computer Physics Communications* **2014**, *185* (2), 604-613.
(9) Henkelman, G.; Uberuaga, B. P.; Jónsson, H. A climbing image nudged elastic band method for finding saddle points and minimum energy paths. *The Journal of Chemical Physics* **2000**, *113* (22), 9901-9904.

(10) Li, C.; Voth, G. A. A quantitative paradigm for water-assisted proton transport through pr oteins and other confined spaces. *Proceedings of the National Academy of Sciences of the United States o f America* **2021**, *118* (49), 1-8.
(11) Díaz Leines, G.; Ensing, B. Path Finding on High-Dimensional Free Energy Landscapes. *Physical Review Letters* **2012**, *109* (2).
